## Supplemental Information for "Modeling the metabolic interplay between a parasitic worm and its bacterial endosymbiont allows the identification of novel drug targets"

### Supplementary information

**Detailed metabolomics methods**

Metabolite extraction – The mass of the weighed worm samples was used to scale the metabolite extraction to a ratio of 16.5 mg / 1 mL extraction solvent. Freezing 80% acetonitrile was added directly to each vial containing the samples, along with zirconium disruption beads (0.5 mm, RPI) and homogenized for 3 min at 4°C in a BeadBlasterTM with a 30 sec on, 30 sec off pattern. The resulting lysate was centrifuged at 21,000 x g for 3 min, and 90% of the supernatant volume was transferred to a 1.5 mL microfuge tube for speed vacuum concentration, no heating. The dry extracts were resolublized in a volume of LCMS grade water 1/10th of that used for the homogenization step, sonicated in a water bath for 3 min, and transferred to a glass insert for analysis.

Data acquisition – All data were acquired by liquid chromatography coupled to high resolution tandem mass spectrometry (LC-MS/MS). The LC column was a MilliporeTM ZIC-pHILIC (2.1 x150 mm, 5 μm) coupled to a Dionex Ultimate 3000TM system and the column oven temperature was set to 25°C for the gradient elution. A flow rate of 100 μL/min was used with the following buffers; A) 10 mM ammonium carbonate in water, pH 9.0, and B) neat acetonitrile. The gradient profile was as follows; 80-20%B (0-30 min), 20-80%B (30-31 min), 80-80%B (31-42 min). Injection volume was set to 2 μL for all analyses (42 min total run time per injection). MS analyses were carried out by coupling the LC system to a Thermo Q Exactive HFTM mass spectrometer operating in heated electrospray ionization mode (HESI). Method duration was 30 min with a polarity switching data-dependent Top 5 method for both positive and negative modes. Spray voltage for both positive and negative modes was 3.5kV and capillary temperature was set to 320°C with a sheath gas rate of 35, aux gas of 10, and max spray current of 100 μA. The full MS scan for both polarities utilized 120,000 resolution with an AGC target of 3e6 and a maximum IT of 100 ms, and the scan range was from 67-1000 m/z. Tandem MS spectra for both positive and negative mode used a resolution of 15,000, AGC target of 1e5, maximum IT of 50 ms, isolation window of 0.4 m/z, isolation offset of 0.1 m/z, fixed first mass of 50 m/z, and 3-way multiplexed normalized collision energies (nCE) of 10, 35, 80. The minimum AGC target was 1e4 with an intensity threshold of 2e5. All data were acquired in profile mode.

Data analysis – Metabolomics data were processed with two approaches called the hybrid analysis and predictive analysis. The resulting relative metabolite intensities for both methods were then processed with an in-house pipeline for statistical analyses and plot generation using a variety of custom Python code and R libraries including: pheatmap, MetaboAnalystR, manhattanly. For the hybrid analysis, peak height intensities were extracted based on the established accurate mass and retention time for each metabolite as adapted from the Whitehead Institute (Chen *et al.*, 2016), and verified with authentic standards and/or high resolution MS/MS manually curated against the NIST14MS/MS (Simón-Manso *et al.*, 2013) and METLIN (Smith *et al.*, 2005) spectral libraries. The theoretical m/z of the metabolite molecular ion was used with a ± 10 ppm mass tolerance window, and a ± 0.2 min peak apex retention time tolerance within the expected elution window (1-2 min). The median mass accuracy vs the theoretical m/z for the library was -4.3 ppm (n=127 detected metabolites). Median retention time range (time between earliest and latest eluting sample for a given metabolite) was 0.24 min (30 min LCMS method). A signal to noise ratio (S/N) of 3X was used compared to blank controls throughout the sequence to report detection, with a floor of 10,000 (arbitrary units). For the predictive analysis, metabolites were detected and quantified by matching the predicted m/z of the metabolite as follows. All predicted model metabolites (n=865) were filtered to the subset with HMDB ID and a neutral chemical formula (n=553). The formula of each metabolite was then used to predict both the positive and negative mode molecular ions excluding adducts, multiply charged species, or more complex ion types i.e., [M+H]+ and [M-H]- ,(n=1106 total m/z values for extraction). A ± 10 ppm mass tolerance window was again used to extract the peak intensities for all m/z values with a ± 0.2 min peak apex retention time tolerance, centering the window on the retention time of the highest intensity peak for each metabolite across all samples. This approach does not discriminate between isomers, for instance 95 of the 553 metabolites in the predicted dataset shared a chemical formula with another metabolite, but in many cases the isomerization was due to stereochemistry (e.g., R-lactic acid vs S-lactic acid) while in other cases formula isomers could confound the data interpretation (dGMP vs AMP). The resulting putative metabolite peak heights were further filtered as follows, after applying the 3X S/N threshold. Metabolites were limited to those that were detected in at least 3 replicates (n=3 per group). Next, an average intensity was calculated for each metabolite across all samples and the data were sorted from high intensity to low intensity. Duplicate metabolite names were removed, thereby keeping the row result from whichever polarity gave a stronger signal (e.g., glutamine was detected in both polarities at 11.5 min, but gave a stronger signal in positive mode).
